# Distance Sampling in R

**DOI:** 10.1101/063891

**Authors:** David L Miller, Eric Rexstad, Len Thomas, Laura Marshall, Jeffrey L Laake

## Abstract

Estimating the abundance and spatial distribution of animal and plant populations is essential for conservation and management. We introduce the `R` package **Distance** that implements distance sampling methods to estimate abundance. We describe how users can obtain estimates of abundance (and density) using the package as well documenting the links it provides with other more specialized `R` packages. We also demonstrate how **Distance** provides a migration pathway from previous software, thereby allowing us to deliver cutting-edge methods to the users more quickly.

## 1. Introduction

Distance sampling (Buckland, Anderson, Burnham, Borchers, and Thomas 2001; Buckland, Anderson, Burnham, Laake, Borchers, and Thomas 2004; Buckland, Rexstad, Marques, and Oedekoven 2015) encompasses a suite of methods used to estimate the density and/or abundance of biological populations. Distance sampling can be thought of as an extension of plot sampling. Plot sampling involves selecting a number of plots (small areas) at random within the study area and counting the objects of interest that are contained within each plot. By selecting the plots at random we can assume that the density of objects in the plots is representative of the study area as a whole. One of the key assumptions of plot sampling is that all objects within each of the plots are counted. Distance sampling relaxes this assumption in that observers are no longer required to detect (i.e., either by eye, video/audio recording, etc.) and count everything within selected plots. While plot sampling techniques are adequate for static populations occurring at high density they are inefficient for more sparsely distributed populations. Distance sampling provides a more efficient solution in such circumstances.

Conventional distance sampling assumes the observer is located either at a point or moving along a line and will observe all objects that occur at the point or on the line. The further away an object is from the point or line (more generally, the sampler or transect) the less likely it is that the observer will see it. We can use the distances to each of the detected objects from the line or point to build a model of the probability of detection given distance from the sampler — the *detection function*. The detection function can be used to infer how many objects were missed and thereby produce estimates of density and/or abundance. Exact distances can be recorded or distances can be collected in bins if exact distances are hard to estimate (sometimes referred to as “grouped” or “interval” data). To ensure that the model is not overly influenced by distances far from zero and that observer time is not spent looking for far away objects, we discard or do not record observations beyond a given *truncation distance* (during analysis or while collecting data in the field).

The Windows program Distance (or “DISTANCE”; for clarity henceforth “Distance for Windows”) can be used to fit detection functions to distance sampling data. It was first released (versions 1.0 - 3.0; principally programmed by Jeff Laake while working at the National Marine Mammal Laboratory) as a console-based application (this in turn was based on earlier software TRANSECT, Burnham, Anderson, and Laake 1980 and algorithms developed in Buckland 1992), before the first graphical interface (Distance for Windows 3.5) was released in November 1998. Since this time it has evolved to include various design and analysis features (Thomas, Buckland, Rexstad, Laake, Strindberg, Hedley, Bishop, Marques, and Burnham 2010). Distance for Windows versions 5 onwards have included `R` (R Core Team 2015) packages as the analysis engines providing additional, more complex analysis options than those offered by the original (`Fortran`) code.

As Distance for Windows becomes increasingly reliant on analyses performed in `R` and many new methods are being developed, we are encouraging the use of our `R` packages directly. `R` provides a huge variety of functionality for data exploration and reproducible research, much more than is possible in Distance for Windows.

Until now those wishing to use our `R` packages for straightforward distance sampling analyses would have had to negotiate the package **mrds** (Laake, Borchers, Thomas, Miller, and Bishop 2015) designed for mark-recapture distance sampling (Burt, Borchers, Jenkins, and Marques 2014), requiring a complex data structure to perform analyses. **Distance** is a wrapper package around **mrds** making it easier to get started with basic distance sampling analyses in `R`. The most basic detection function estimation only requires a numeric vector of distances. Here we demonstrate how to use **Distance** to fit detection functions, perform model checking and selection, and estimate abundance.

### 1.1. Distance sampling

The distribution of the observed distances is a product of detectability (sometimes referred to as “perception bias”; Marsh and Sinclair 1989) and the distribution of the animals with respect to the line or point. Our survey design allows us to assume a distribution for the animals with respect to the sampler.

For line transect studies we assume that objects are uniformly distributed with respect to the line (i.e., the number of animals available for detection is the same at all distances). For point transect surveys, area increases linearly with radial distance, implying a triangular distribution with respect to the point. Figure 1 shows how these distributions, when combined with a detection function, give rise to the observed distribution of recorded distances.

**Figure. 1.**
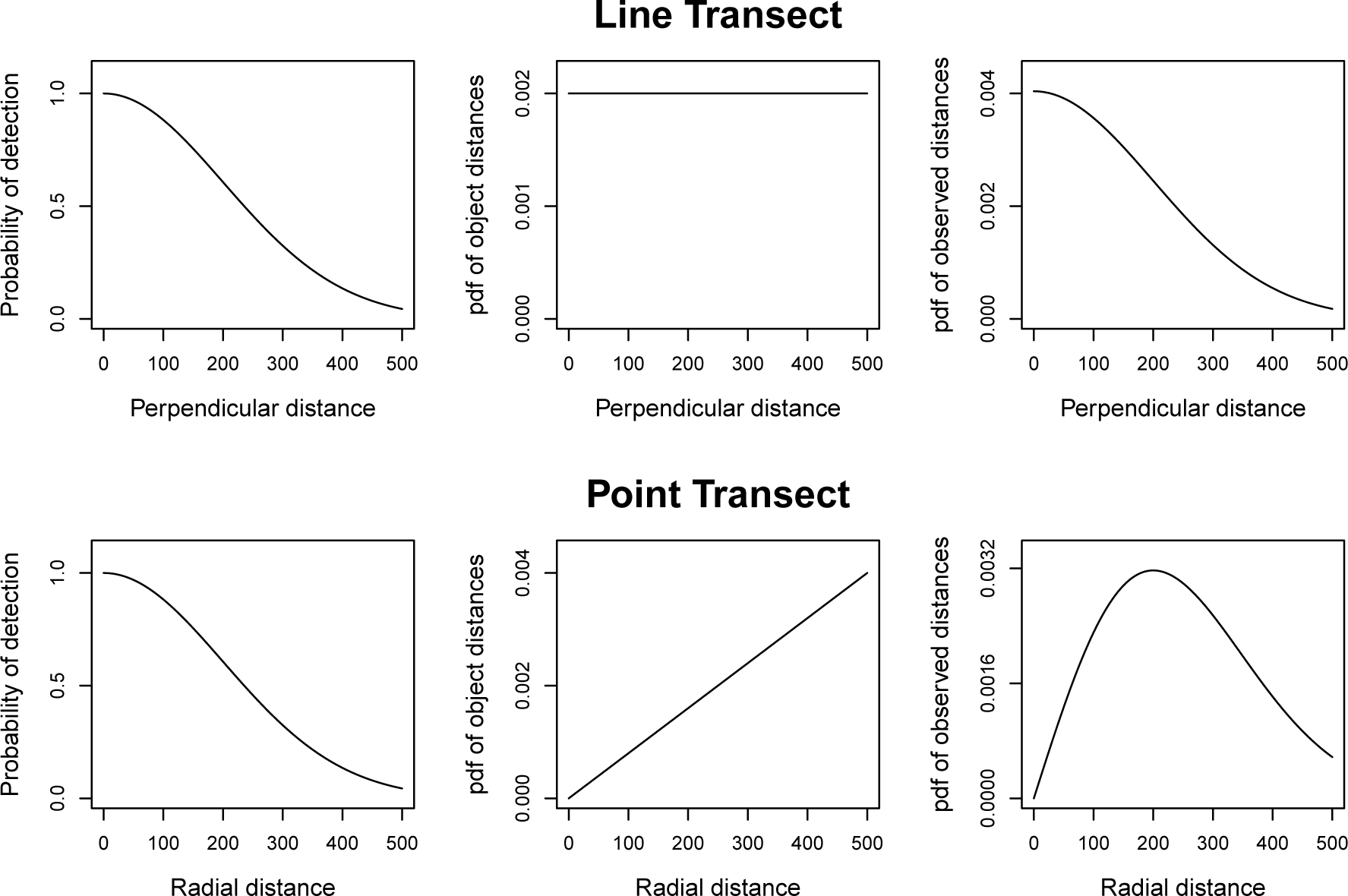
Panels show an example detection function (left), the probability density function of object distances (middle) and the resulting pdf of detection distances (right) for line transects (top row) and point transects (bottom row). The pdf of observed detection distances in the right hand plots are obtained by multiplying the detection function by the pdf of object distances and rescaling. In this example, detection probability becomes effectively zero at 500 (distances shown on *x*-axis are arbitrary).

Figure 2 shows simulated sampling of a population of 500 objects using line and point transects and their corresponding histograms of observed detection distances. Note that for the purposes of distance sampling an “object” may either refer to an individual in a population or a cluster (or group) of individuals.

**Figure. 2.**
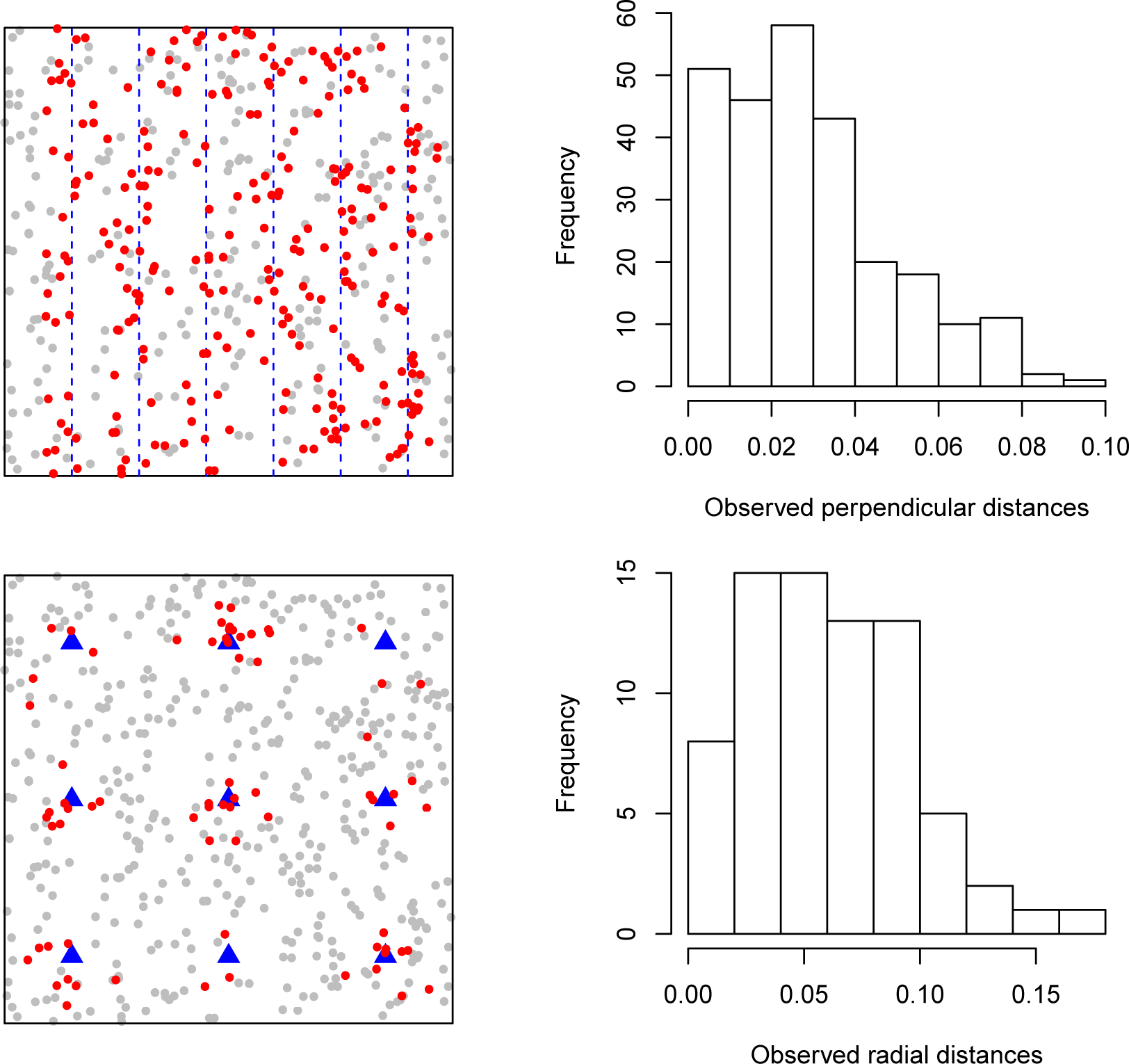
Examples of line (top row) and point (bottom row) transect sampling. Left side plots show an example of a survey of the unit square containing a population of 500 objects; blue dashed lines (top plot) and triangles (bottom plot) indicate sampler placement, red dots indicate detected individuals and grey dots give the locations of unobserved individuals. The right side of the figure shows histograms of observed distances.

Good survey design is essential to produce reliable density and abundance estimates from distance sampling data. Survey design is beyond the scope of this article but we refer readers to (Buckland *et al*. 2001, Chapter 7) and (Buckland *et al*. 2015, Chapter 2) for introductory information; Strindberg, Buckland, and Thomas (2004) contains information on automated survey design using geographical information systems (GIS); Thomas, Williams, and Sandilands (2007) gives an example of designing a distance sampling survey in a complex region. We also note that Distance for Windows includes a GIS survey design tool (Thomas *et al*. 2010).

**Distance** provides a selection of candidate functions to describe the probability of detection and estimates the associated parameters using maximum likelihood estimation. The probability of detecting an object may not only depend on how far it is from the sampler but also on other factors such as weather conditions, ground cover, cluster size etc (Marques, Thomas, Fancy, and Buckland 2007). The **Distance** package also allows the incorporation of such covariates into the detection function allowing the detection function scale parameter to vary based on these covariates.

Having estimated the detection function’s parameters, one can then integrate out distance from the function (as the detection function describes the probability of detection *given* distance) to get an “average” probability of detection (in the sense of averaging over distances, conditional on any observed covariates), which can be used to correct the observed counts. Summing the corrected counts gives an estimate of abundance in the area covered by surveys, which can be multiplied up to the total study area.

In addition to randomly placed samplers distance sampling relies on three other main assumptions. Firstly, all objects directly on the transect line or point (i.e., those at zero distance) are detected (see “Extensions” for methods to deal with the situation when this is not possible). Secondly, objects are stationary or detected prior to any movement. Thirdly, distance to the object must be measured accurately, or the observation allocated to the correct distance bin for grouped data. Depending on the survey species some of these assumptions may be more difficult to meet than others. Further information on field methods to help meet these assumptions can be found in Buckland *et al*. (2001) and Buckland *et al*. (2015).

The rest of the paper has the following structure: we describe data formatting for **Distance**; candidate detection function models are described and examples fitted in `R`. We then show how to perform model checking, goodness of fit testing and model selection. We go on to show how to estimate abundance, including stratified estimates of abundance. The final two sections of the article look at extensions (both in terms of methodology and software) and put the package in a broader context amongst other `R` packages used for estimating the abundance of biological populations from distance sampling data.

## 2. Data

We introduce two example analyses performed in **Distance**: one line transect and one point transect. These data sets have been chosen as they represent typical data seen in practice. The below example analyses are not intended to serve as guidelines, but to demonstrate features of the software. Practical advice on approaches to analysis is given in Thomas *et al*. (2010).

### 2.1. Minke whales

The line transect data have been simulated from models fitted to Antarctic minke whale (*Balaenoptera bonaerensis*) data. These data were collected as part of the International Whaling Commission’s International Decade of Cetacean Research Southern Ocean Whale and Ecosystem Research (IWC IDCR-SOWER) programme 1992-1993 austral summer surveys. Data consist of 99 observations on 25 transects, which were stratified based on location (near or distant from ice edge) and effort data (transect lengths). Further details on the survey are available in Branch and Butterworth (2001) (data are simulated based on the design used for “1992/93 Area III” therein).

### 2.2. Amakihi

The point transect data set consists of 1485 observations of Amakihi (*Hemignathus virens*; a Hawaiian songbird), collected at 41 points between 1992 and 1995. The data include distances and two covariates collected during the survey: observer (a three level factor), time after sunrise (transformed to minutes (continuous) or hours (factor) covariates). Data are analysed comprehensively in Marques *et al*. (2007).

### 2.3. Data setup

Generally, data collected in the field will require some formatting before use with **Distance**, though there are a range of possible formats, dependent on the model specification and the output required:

- In the simplest case, where the objective is to estimate a detection function and exact distances are collected, all that is required is a numeric vector of distances.
- To include additional covariates into the detection function (see “Detection functions”) a `data.frame` is required. Each row of the `data.frame` contains the data on one observation. The `data.frame` must contain a column named `distance` (containing the observed distances) and additional named columns for any covariates that may affect detectability (for example `observer` or `seastate`). The column name `size` is reserved for the cluster sizes (sometimes referred to as group sizes) in this case each row represents an observation of a cluster rather than individual (see Buckland *et al*. 2001, Section 1.4.3 for more on defining clusters and “Extensions” below for one approach to dealing with uncertain cluster size). Additional reserved names include `object` and `detected`, these are not required for conventional distance sampling and should be avoided (see “Extensions” for an explanation of their use).
- To estimate density or to estimate abundance beyond the sampled area, additional information is required. Additional columns should be included in the `data.frame` specifying: `Sample.Label`, the ID of the transect; `Effort`, transect effort (for lines their length and for points the number of times that point was visited); `Region.Label`, the stratum containing the transect (which may be from pre- or post-survey stratification, see “Estimating abundance and variance”); `Area`, the area of the strata. Transects which were surveyed but have no observations must be included in the data set with `NA` for `distance` and any other covariates. We refer to this data format (where all information is contained in one table) as “flatfile” as it can be easily created in a single spreadsheet.

As we will see in “Extensions”, further information is also required for fitting more complex models.

If distances were collected in bins the column `distance` is replaced by two columns `distbegin` and `distend` referring to the distance interval start and end cutpoints. More information on binned data is included in Buckland *et al*. (2001) sections 4.5 and 7.4.1.2.

The columns `distance, Area` and (in the case of line transects) `Effort` have associated units (though these are not explicitly included in a **Distance** analysis). We recommend that data in these columns are converted to SI units before starting any analysis to ensure that resulting abundance and density estimates have sensible units. For example, if distances from a line transect survey are recorded in metres, the `Effort` columns should contain line lengths also in metres and the `Area` column gives the stratum areas in square metres. This would lead to density estimates of animals per square metre.

The minke whale data follows the “flatfile” format given in the last bullet point:

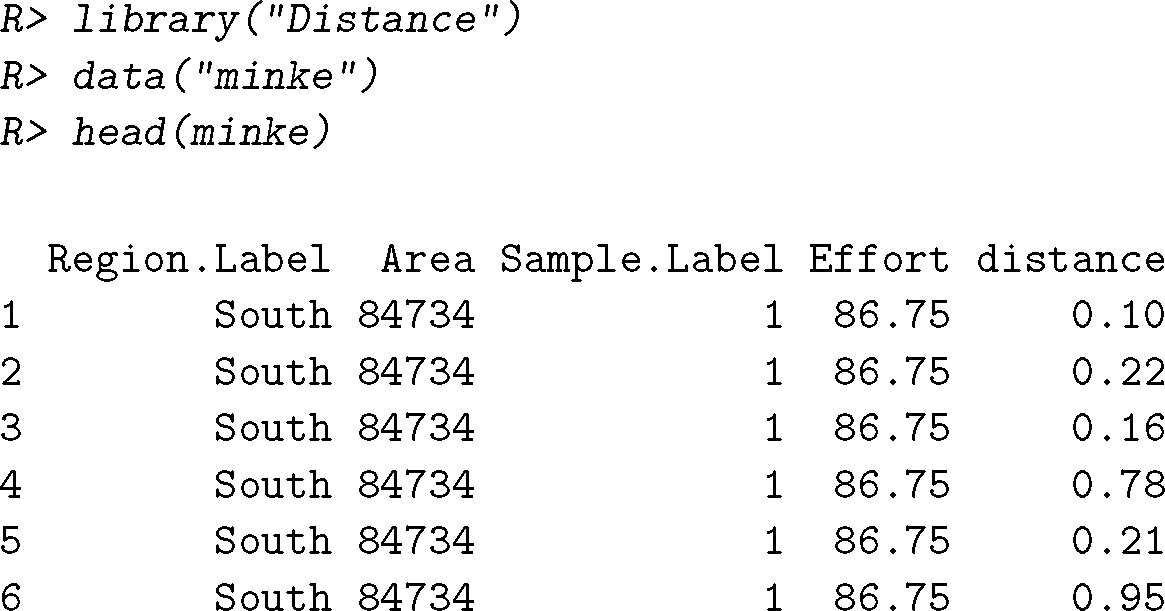

Whereas the amakihi data lacks effort and stratum data:

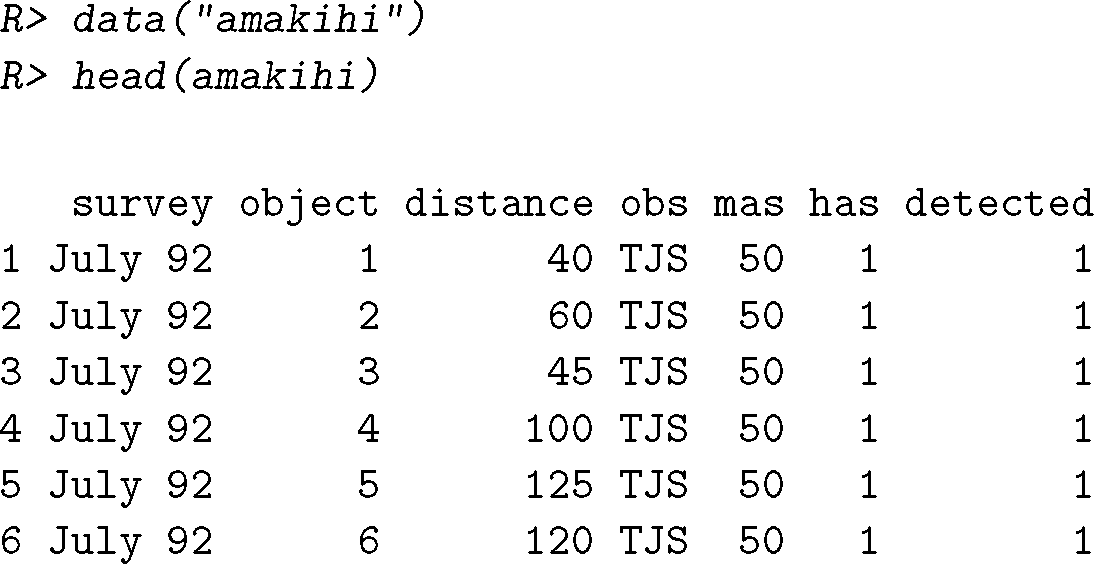

We will explore the consequences of including effort and stratum data in the analysis below.

## 3. Detection functions

The detection function models the probability `P`(object detected | object at distance y) and is usually denoted *g*(*y*; ***θ***) where *y* is distance (from a line or point) and ***θ*** is a vector of parameters to be estimated. Our goal is to estimate an *average probability of detection* (*p*, average in the sense of an average over distance from 0 to truncation distance *w*), so we must integrate out distance from the detection function. Letting *x* denote a perpendicular distance from a line and *r* denote radial distance from a point:

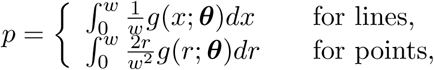

where the fractions pre-multiplying the detection function describe the distribution of objects with respect to the sampler, taking into account the geometry of the sampler (usually referred to as the *probability density function of (object) distances* and denoted *π*(*y*); Buckland *et al*. 2001, Chapter 3). Figure 1 shows the relationship between the detection function, the probability density function of object distances and the probability density function of observed distances.

Models for the detection function are expected to have the following properties (Buckland *et al*. 2015, Chapter 5):

- *Shoulder*: we expect observers to be able to see objects near them, not just those directly in front of them. We therefore expect the detection function to be flat near the line or point.
- *Non-increasing*: we do not think that observers should be more likely to see distant objects than those nearer the transect. If this occurs, it usually indicates an issue with survey design or field procedures (for example that the distribution of objects with respect to the line, *π*(*y*), is not what we expect), so we do not want the detection function to model this (Marques, Buckland, Tosh, McDonald, and Borchers 2010; Marques, Buckland, Bispo, and Howland 2012; Miller and Thomas 2015).
- *Model robust*: models should be flexible enough to fit many different shapes.
- *Pooling robust*: many factors can affect the probability of detection and it is not possible to measure all of these. We would like models to produce unbiased results without inclusion of these factors.
- *Estimator efficiency*: we would like models to have low variances, given they satisfy the other properties above (which, if satisfied, would give low bias).

The shoulder condition also implies that it is crucial that the detection function accurately models detectability at small distances and that we are less worried by its behaviour further away from 0. Given these criteria, we can formulate models for *g*.

### 3.1. Formulations

There is a wide literature on possible formulations for the detection function (Buckland 1992; Eidous 2005; Becker and Quang 2009; Giammarino and Quatto 2014; Miller and Thomas 2015; Becker and Christ 2015). **Distance** includes the most popular of these models. Here we detail the most popular detection function approach: “key function plus adjustments” (K+A).

*Key function plus adjustments (K+A)*

Key function plus adjustment terms (or adjustment series) models are formulated by taking a “key” function and optionally including “adjustments” to it to improve the fit (Buckland 1992). Mathematically we formulate this as:

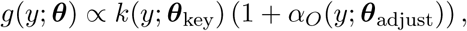

where *k* is the key function and *α_O_* is some series of functions (given in Table 1), described as an *adjustment of order O*. Subscripts on the parameter vector indicate those parameters belonging to each part of the model (i.e., ***θ*** = (***θ***_key_, ***θ***_adjust_)).

Available models for the key are as follows:

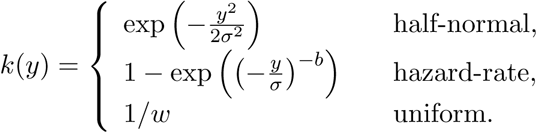

**Table 1.**
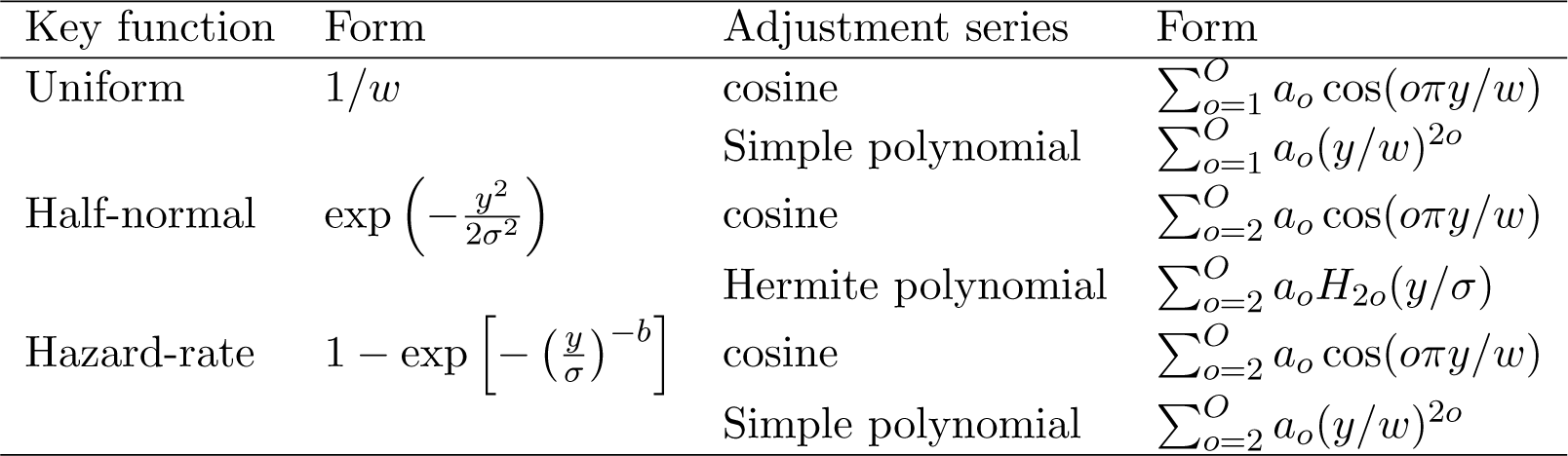
Modelling options for key plus adjustment series models for the detection function. For each key function the default adjustments are cosine in **Distance**. Note that in the adjustments functions distance is divided by the width or the scale parameter to ensure the shape of adjustment independent of the units of *y* (Marques *et al*. 2007); defaults are shown here, though either can be selected to rescale the distances.

Possible modelling options for key and adjustments are given in Table 1 and illustrated in Figures 3 and 4. We select the number of adjustment terms (*K*) by AIC (further details in “Model checking and model selection”).

When adjustment terms are used it is necessary to standardise the results to ensure that *g*(0) = 1:

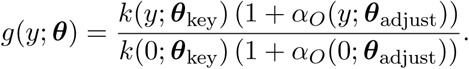

A disadvantage of K+A models is that we must resort to constrained optimisation (via the **Rsolnp** package; Ghalanos and Theussl 2014) to ensure that the resulting detection function is monotonic non-increasing over its range.

It is not always necessary to include adjustments (except in the case of the uniform key) and in such cases we refer to these as “key only” models (see the next section and “Model checking and model selection”). Adjustment terms increase the flexibility of the detection function but this added flexibility comes at the expense of additional parameters to be estimated. The analyst must consider whether the process giving rise to the distribution of detection distances should be modelled with shapes depicted in Figure 4.

**Figure. 3.**
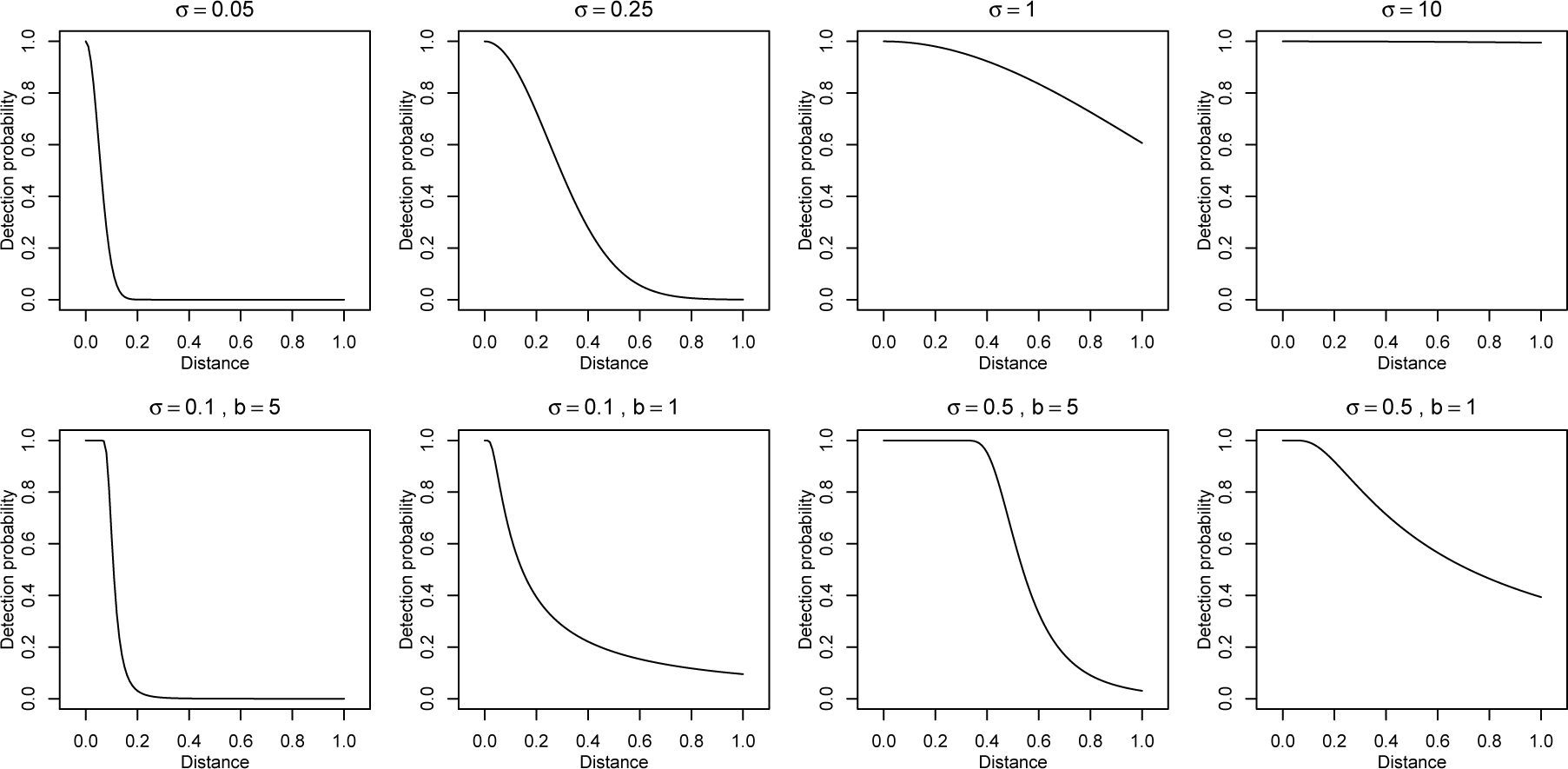
Half-normal (top row) and hazard-rate (bottom row) detection functions without adjustments, varying scale (*σ*) and (for hazard-rate) shape (*b*) parameters (values are given above the plots). On the top row from left to right, the study species becomes more detectable (higher probability of detection at larger distances). The bottom row shows the hazard-rate model’s more pronounced shoulder.

**Figure. 4.**
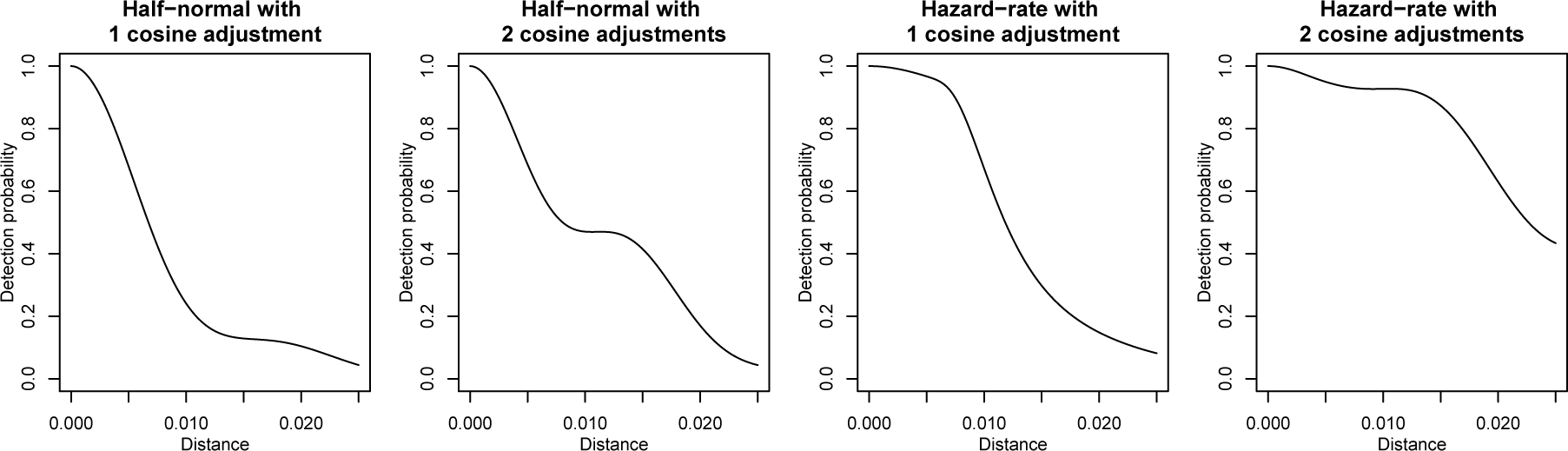
Possible shapes for the detection function when cosine adjustments are included for half-normal and hazard-rate models.

*Covariates*

There are many factors that can affect the probability of detecting an object: observer skill, cluster size (if objects occur in clusters), the vessel or platform used, sea state, other weather conditions, time of day and more. In **Distance** we assume that these covariates affect detection only via the scale of the detection function (and do not affect the shape).

Covariates can be included in this formulation by considering the scale parameter from the half-normal or hazard-rate detection functions as a(n exponentiated) linear model of the (*J*) covariates (**z**; a vector of length *J* for each observation):

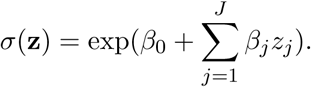

Including covariates has an important implication for our calculation of detectability. We do not know the true distribution of the covariates, we therefore must either: (*i*) model the distribution of the covariates and integrate the covariates out of the joint density (thus making strong assumptions about their distribution), or (*ii*) calculate the probability of detection conditional on the observed values of the covariates (Marques and Buckland 2003). We opt for the latter:

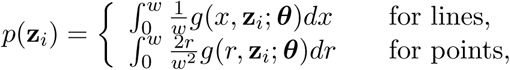

where ***z**_i_* is the vector of *J* covariates associated with observation *i*. For covariate models, we calculate a value of “average” probability of detection (average in the sense of distance being integrated out) per observation. There are as many unique values of *p*(***z**_i_*) as there are unique covariate combinations in our data.

K+A models that include covariates and one or more adjustments cannot be guaranteed to be monotonic non-increasing for all covariate combinations. Without a model for the distribution of the covariates, it is not possible to know what the behaviour of the detection function will be across the ranges of the covariates. As such we cannot set meaningful constraints on monotonicity. For this reason, we advise caution when using both adjustments and covariates in a detection function (see Miller and Thomas 2015, for an example of when this can be problematic and an alternative detection function formulation to solve this issue).

### 3.2. Fitting detection functions in R

A detection function can be fitted in **Distance** using the `ds` function. Here we apply some of the possible formulations for the detection function we have seen above to the minke whale and amakihi data.

*Minke whale*

First we fit a model to the minke whale data, setting the truncation at 1.5km and using the default options in **ds** very simply:

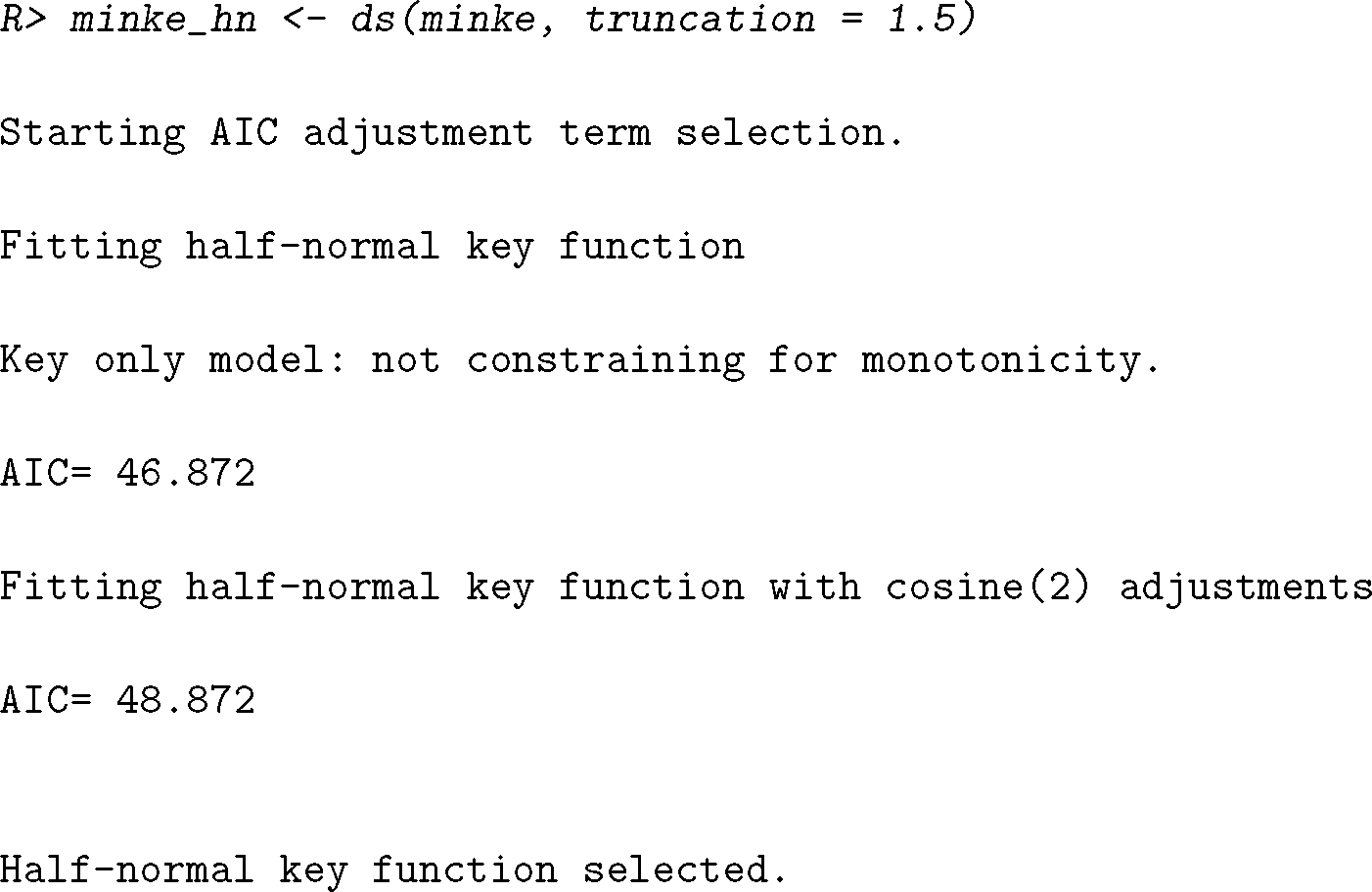

Note that when there are no covariates in the model, `ds` will add adjustment terms to the model until there is no improvement in AIC.

Figure 5 (left panel) shows the result of calling `plot` on the resulting model object. We can also call `summary` on the model object to get summary information about the fitted model (we postpone this until the next section).

A different form for the detection function can be specified via the `key=` argument to `ds`. For example, a hazard rate model can be fitted as:

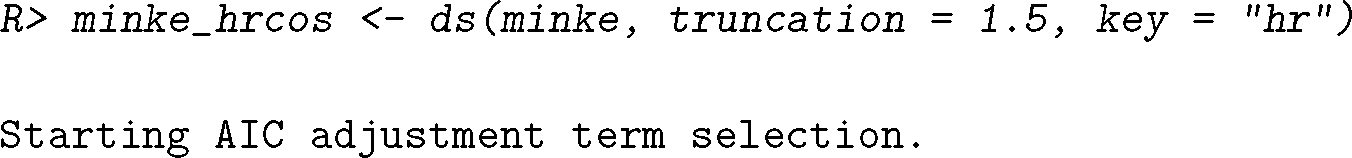
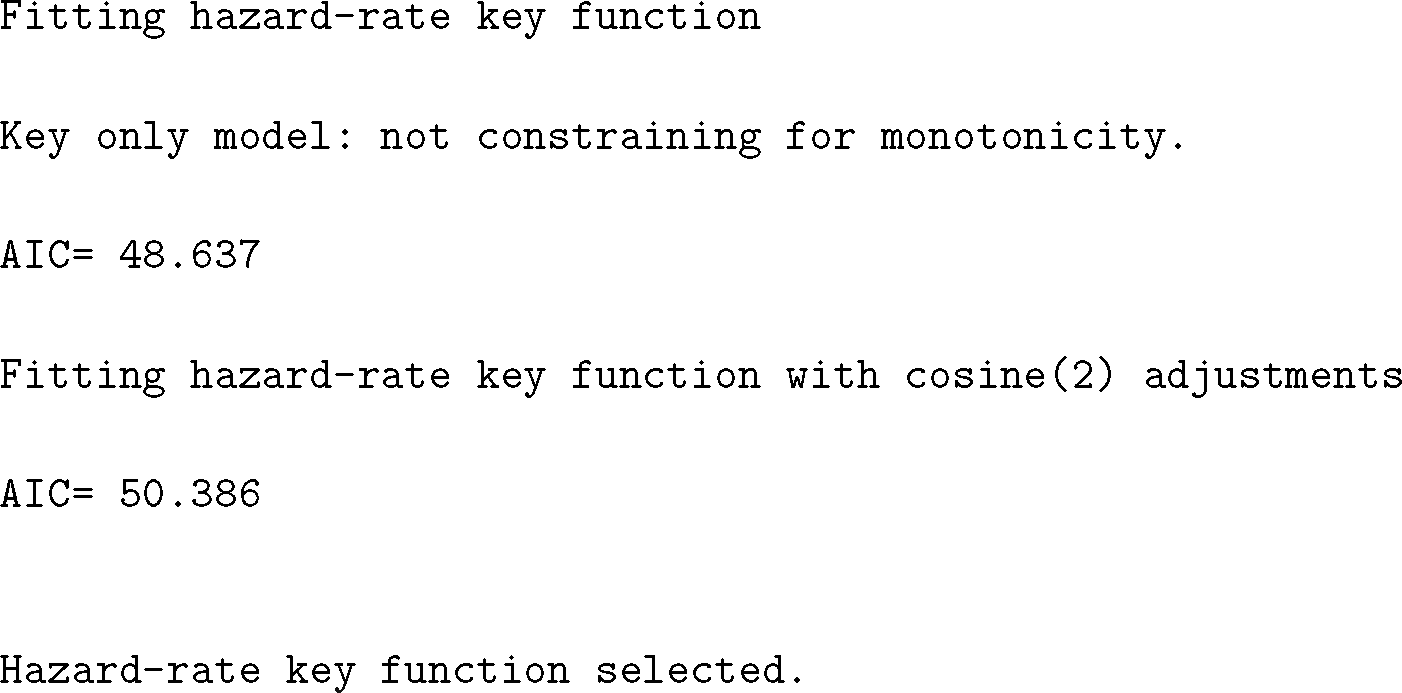

Here `ds` also fits the hazard-rate model then hazard-rate with a cosine adjustment but the AIC improvement is insufficient to select the adjustment, so the hazard-rate key-only model is returned.

Other adjustment series can be selected using the `adjustment=` argument and specific orders of adjustments can be set using `order=`. For example, to specify a uniform model with cosine adjustments of order 1 and 2 we can write:

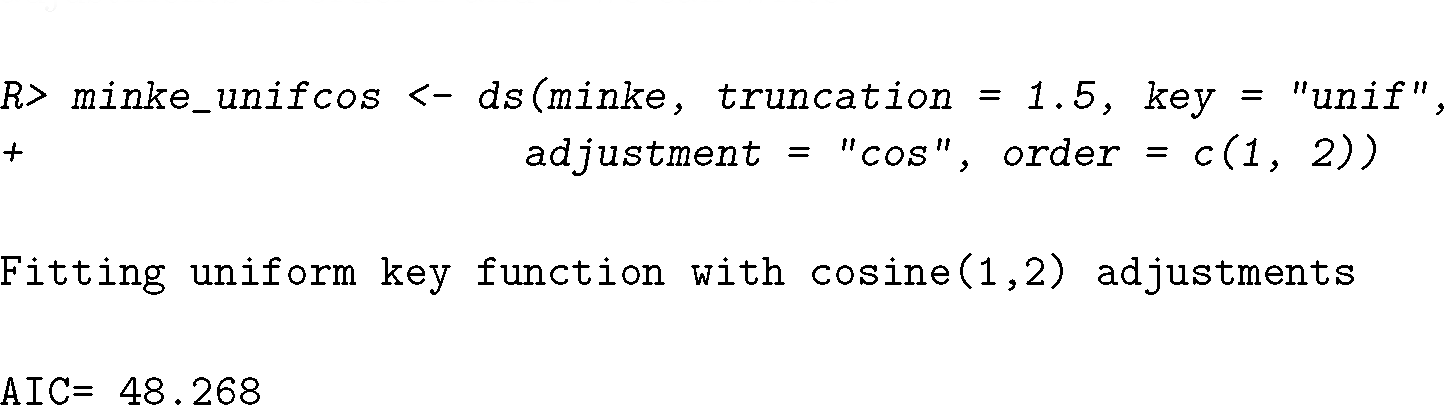

Hermite polynomial adjustments use the code •`herm`• and simple polynomials •`poly`•, adjustment order should be in line with Table 1.

*Amakihi*

`ds` assumes the data given to it has been collected as line transects, but we can switch to point transects using the argument `transect=•point•`. We can include covariates in the scale parameter via the `formula=∼…` argument to `ds`. A hazard-rate model for the amakihi that includes observer as a covariate and a truncation distance of 82.5m (Marques *et al*. 2007) can be specified using :

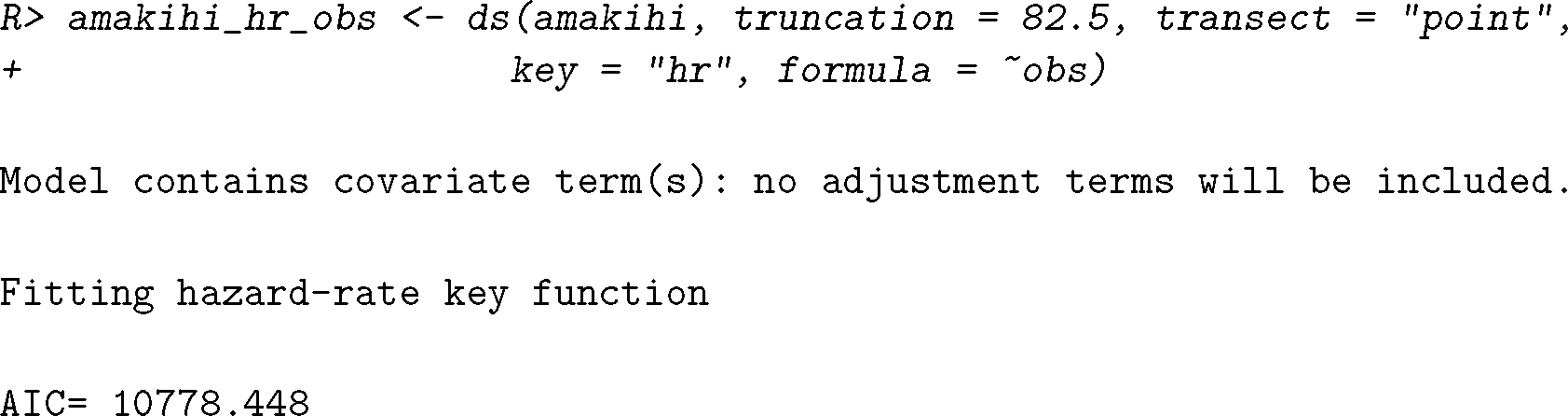
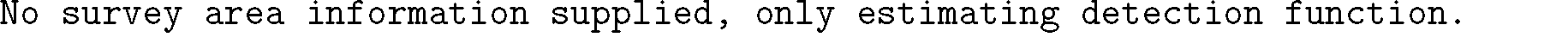

Note that here, unlike with the minke whale data, `ds` warns us that we have only supplied enough information to estimate the detection function (not density or abundance).

While automatic AIC selection is performed on adjustment terms, model selection for covariates must be performed manually. Here we add a second covariate: minutes after sunrise. We will compare these two models further in the following section.

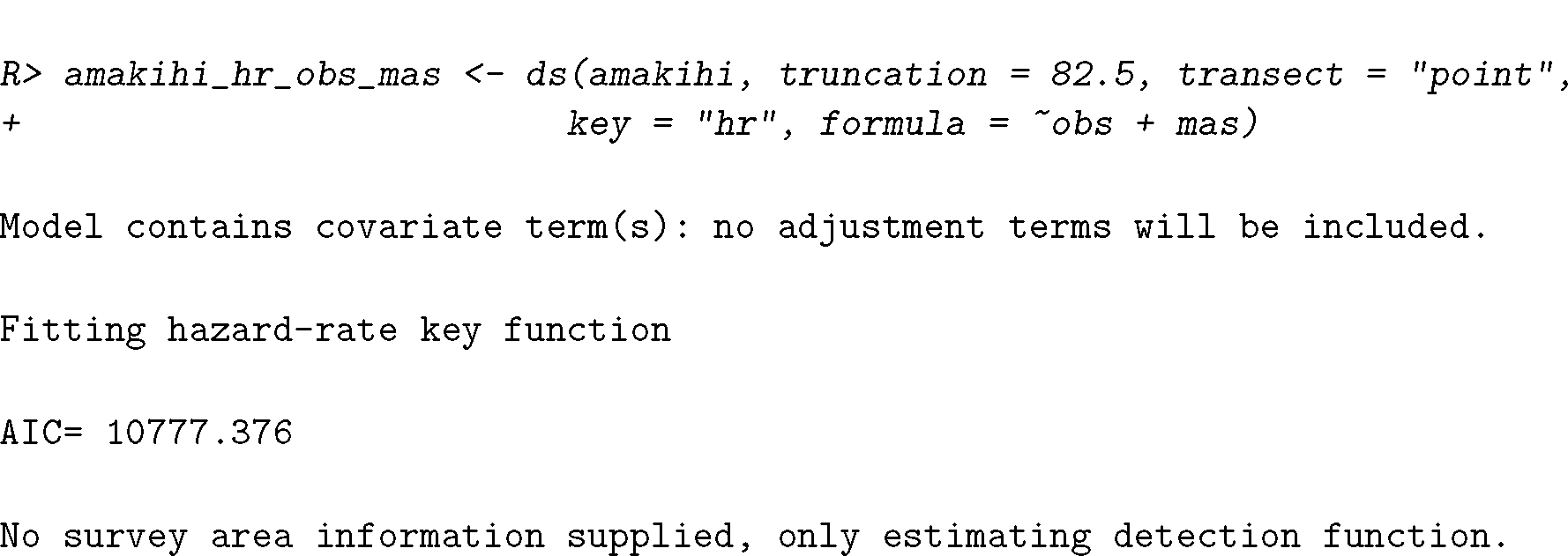

As with the minke whale model, we can plot the resulting models (Figure 5, middle and right panels). However, for point transect studies, probability density function plots give a better sense of model fit than the detection function plots. This is because when plotting the detection function for point transect data, the histogram must be rescaled to account for the geometry of the point sampler. The amakihi models included covariates, so the plots show the detection function averaged over levels/values of the covariate. Points on the plot indicate probability of detection for each observation. For the `amakihi_hr_obs` model we see fairly clear levels of the observer covariate in the points. Looking at the right panel of Figure 5, this is less clear when adding minutes after sunrise (a continuous covariates) to the model.

**Figure. 5.**
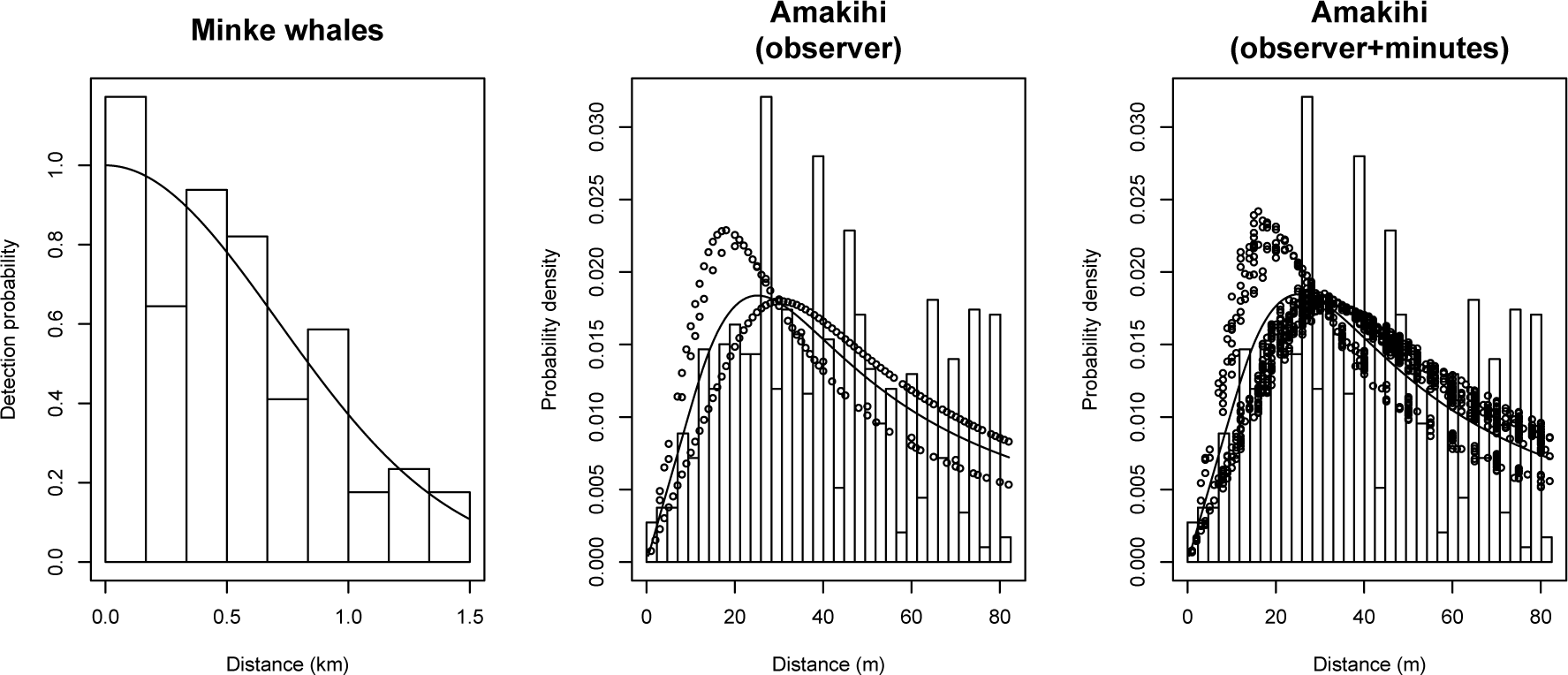
Left: fitted detection function overlayed on the histogram of observed distances for the minke whale data using half-normal model. Centre and right: plots of the probability density function for the amakihi models. Centre, hazard-rate with observer as a covariate; right, hazard-rate model with observer and minutes after sunrise as covariates. Points indicate probability of detection for a given observation (given that observation’s covariate values) and lines indicate the average detection function (averaged over covariates, observer or observer+minutes after sunrise).

## 4. Model checking and model selection

As with models fitted using `lm` or `glm` in `R`, we can use `summary` to give useful information about our fitted model. For our hazard-rate model of the amakihi data, with observer as a covariate:

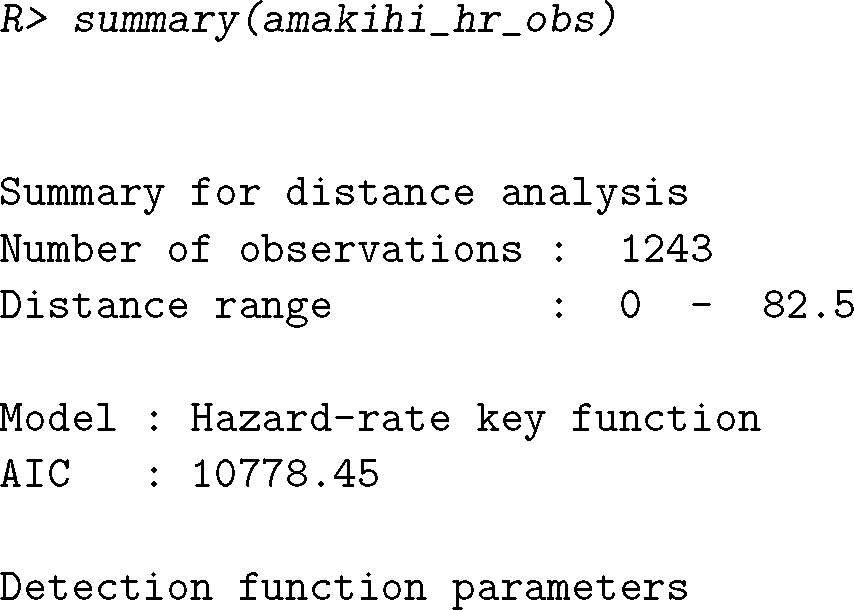

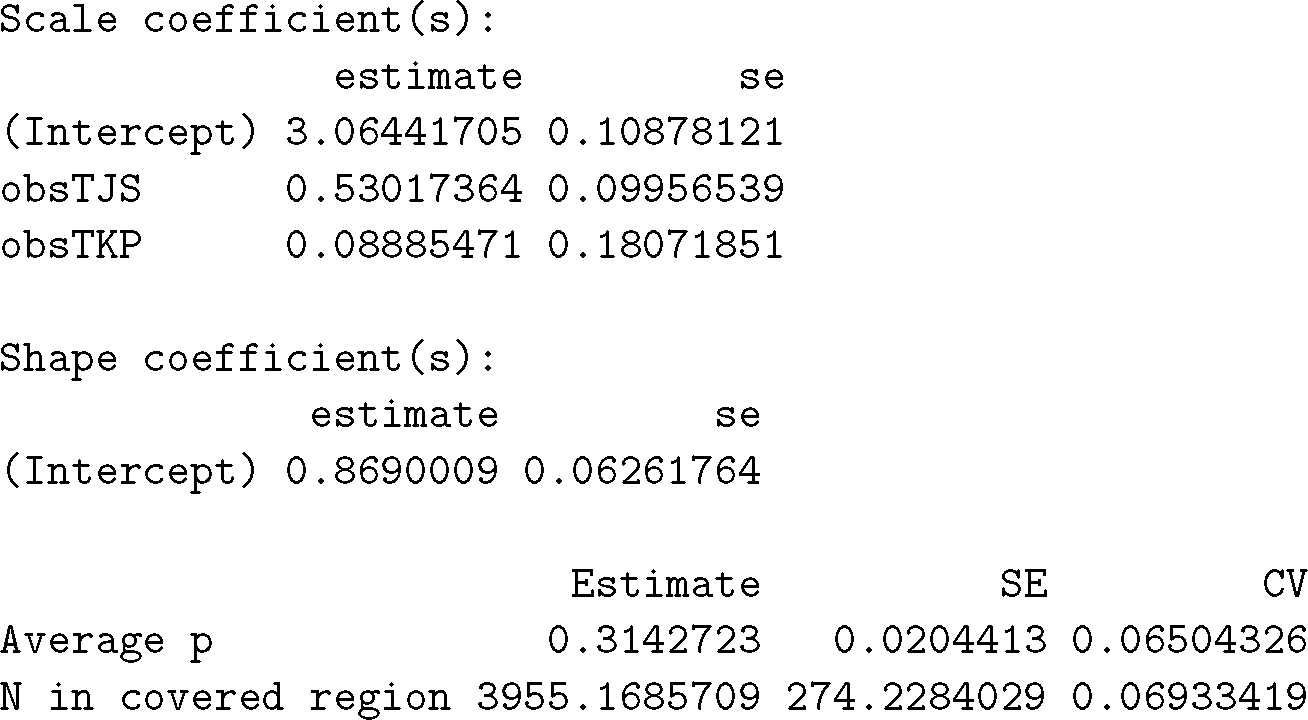

This summary information includes details of the data and model specification, as well as the values of the coefficients (*β_j_*) and their uncertainties, an “average” value for the detectability (see “Estimating abundance and variance” for details on how this is calculated) and its uncertainty. The final line gives an estimate of abundance in the area covered by the survey (see the next section; though note this estimate does not take into account cluster size).

### 4.1. Goodness of fit

We use a quantile-quantile plot (Q-Q plot) to visually assess how well a detection functions fits the data when we have exact distances. The Q-Q plot compares the cumulative distribution function (CDF) of the fitted detection function to the distribution of the data (empirical distribution function or EDF). The Q-Q plots in **Distance** plot a point for every observation. The EDF is the proportion of points that have been observed at a distance equal to or less than the distance of that observation. The CDF is calculated from the fitted detection function as the probability of observing an object at a distance less than or equal to that of the given observation. This can be interpreted as assessing whether the number of observations up to a given distance is in line with what the model says they should be. As usual for Q-Q plots, “good” models will have values close to the line *y* = *x*, poor models will show greater deviations from that line.

Q-Q plots can be inspected visually, though this is prone to subjective judgments. Therefore, we also quantify the Q-Q plot’s information using a Cramér-von Mises test (Burnham, Buckland, Laake, Borchers, Bishop, and Thomas 2004) to test whether points from the EDF and CDF are from the same distribution. The Cramér-von Mises test uses the sum of all the distances between a point on the Q-Q plot and the line *y* = *x* to form a test statistic. As it takes into account more information and is therefore more powerful, the Cramér-von Mises is generally more powerful than the Kolmogorov-Smirnov test, which uses the largest difference between a point on the Q-Q plot and the line *y* = *x* (which is also available in **Distance**, though is not produced by default as it requires computationally demanding bootstraps). A significant result from either test gives evidence against the null hypothesis (that the data arose from the fitted model), suggesting that the model does not fit the data well.

We can generate a Q-Q plot and test results using the `gof_ds` function. Figure 6 shows the goodness of fit tests for two models for the amakihi data. We first fit a half-normal model without covariates or adjustments (setting `adjustment=NULL` will force `ds` to fit a model with no adjustments):

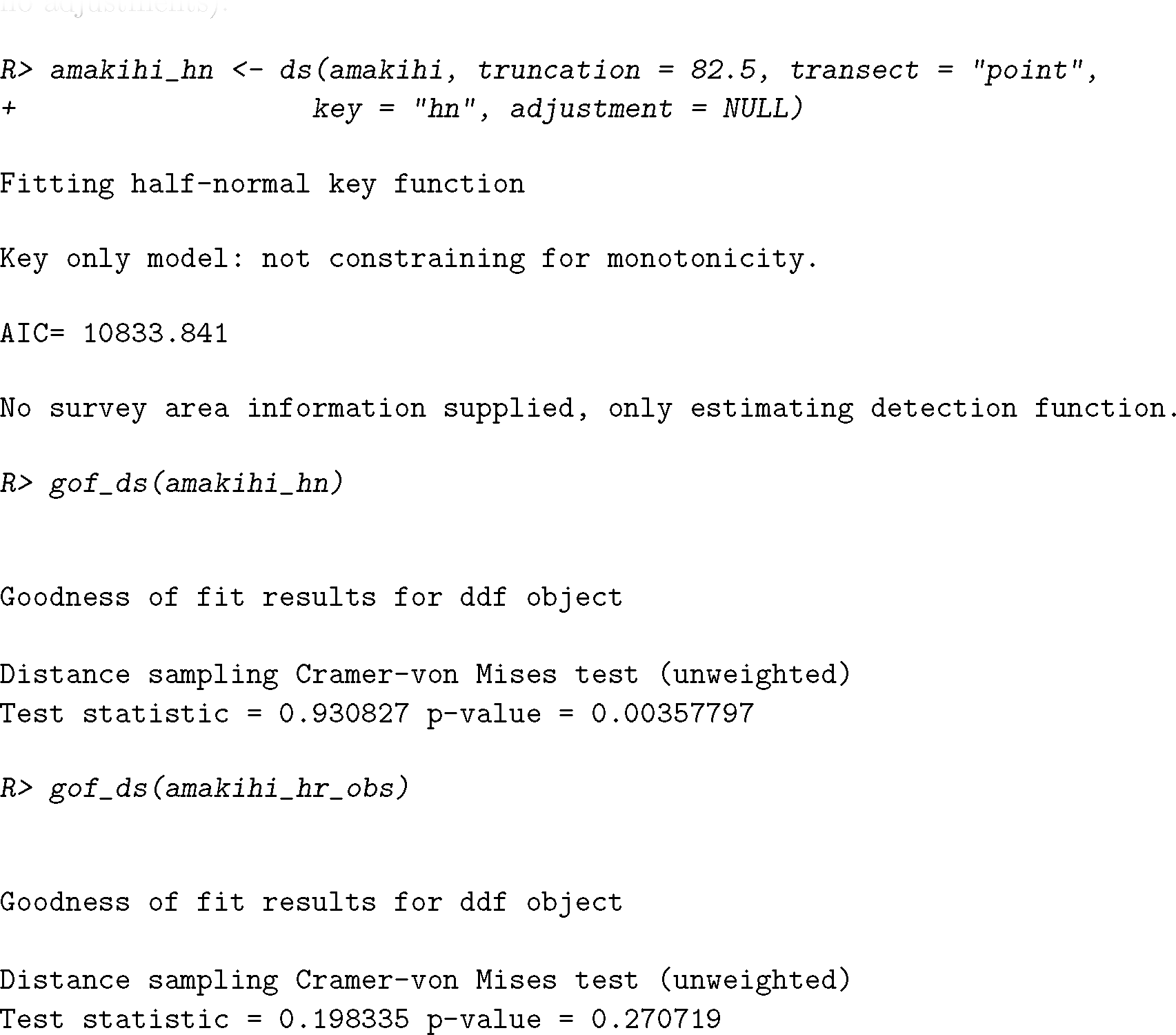

**Figure. 6.**
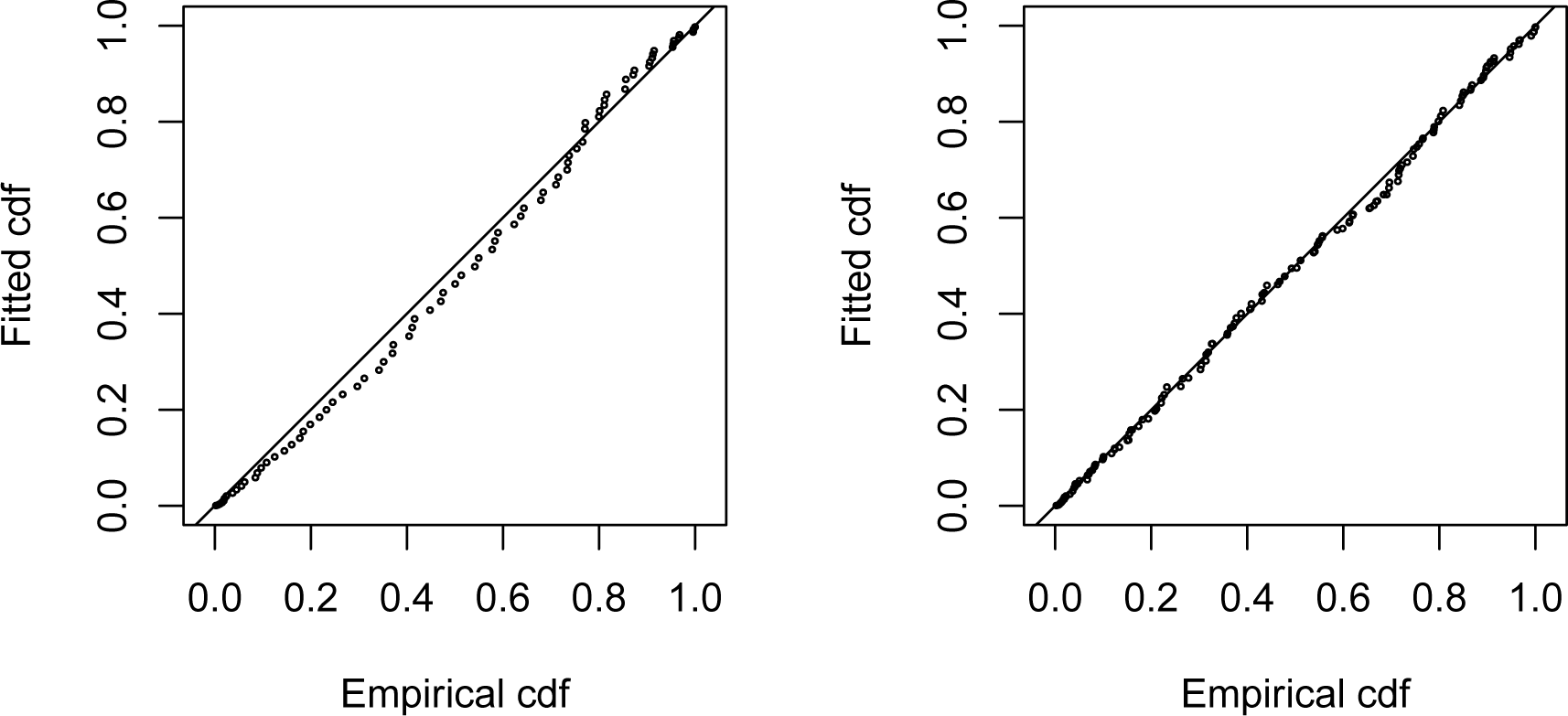
Comparison of quantile-quantile plots for a half-normal model (no adjustments, no
covariates; left) and hazard-rate model with observer as a covariate (right) for the amakihi
data.

We conclude that the half-normal model should be discarded. Both hazard-rate models (output only shown for hazard-rate model with observer and minutes after sunrise) had nonsignificant goodness of fit test statistics and are both therefore deemed plausible models. The corresponding Q-Q plots are shown in Figure 6, comparing the half-normal model with the hazard-rate model with observer and minutes after sunrise included.

For non-exact distance data, a *χ*^2^-test can be used to assess goodness of fit (see Buckland *et al*. 2001, Section 3.4.4). *χ*^2^-test results are produced by `gof_ds` when binned/grouped distances are provided.

### 4.2. Model selection

Once we have a set of plausible models, we can use Akaike’s Information Criterion (AIC) to select between models (see e.g. Burnham and Anderson 2003). **Distance** includes a function to create table of summary information for fitted models, making it easy to get an overview of a large number of models. The `summarize_ds_models` function takes models as input and can be especially useful when paired with **knitr**’s `kable` function to create summary tables for publication (Xie 2015). An example of this output (with all models included) is shown in Table 2 and was generated by the following call to `summarize_ds_models`:

**Table 2.** Summary for the detection function models fitted to the amakihi data. “C-vM” stands for Cramér-von Mises, *P_a_* is average detectability (see “Estimating abundance and variance”), se is standard error. Models are sorted according to AIC.

| Model | Key function | Formula | C-vM p-value | $\hat{P}_a$ | $se(\hat{P}_a)$ | $\Delta AIC$ |
| --- | --- | --- | --- | --- | --- | --- |
| `amakihi_hr_obs_mas` | Hazard-rate | `obs + mas` | 0.389 | 0.319 | 0.020 | 0.000 |
| `amakihi_hr_obs` | Hazard-rate | `obs` | 0.271 | 0.314 | 0.020 | 1.073 |
| `amakihi_hn` | Half-normal | `1` | 0.004 | 0.351 | 0.011 | 56.465 |

`summarize_ds_models(amakihi_hn, amakihi_hr_obs, amakihi_hr_obs_mas)`

In this case we may be skeptical about the top model as selected by AIC being truly better than the second best model, as there is only a very small difference in AICs. Generally, if the difference between AICs is less than 2, we may investigate multiple “best” models, potentially resorting to the simplest of these models. In the authors’ experience, it is often the case that models with similar AICs will have similar estimates probabilities of detection, so in practice there is little difference in selecting between these models. It is important to note that comparing AICs between models with different truncations is not appropriate, as models with different truncation use different data.

## 5. Estimating abundance and variance

Though fitting the detection function is the primary modelling step in distance sampling, we are really interested in estimating density or abundance. It is also important to calculate our uncertainty for these estimates. This section addresses these issues mathematically before showing how to estimate abundance and its variance in `R`.

### 5.1. Abundance

We wish to obtain the abundance in a study region, of which we have sampled a random subset. To do this we first calculate the abundance in the area we have surveyed (the *covered area*) to obtain 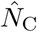, we can then scale this up to the full study area by multiplying it by the ratio of covered area to study area. We discuss other methods for spatially explicit abundance estimation in “Extensions”.

First, to estimate abundance in the covered area (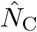), we use the estimates of detection probability (the 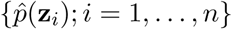, above) in a Horvitz-Thompson-like estimator:

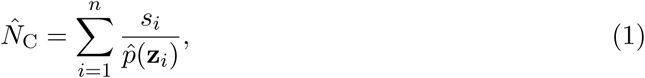

where *s_i_* are the sizes of the observed clusters of objects, which are all equal to 1 if objects only occur singly (Borchers and Burnham 2004). Thompson (2002) is the canonical reference to this type of estimator. Intuitively, we can think of the estimates of detectability 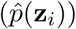 as “inflating” the cluster sizes (*s_i_*), we then sum over the detections (*i*) to obtain the abundance estimate. For models that do not include covariates, 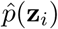 is equal for all *i*, so this is equivalent to summing the clusters and inflating that sum by dividing through by the corresponding 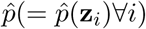.

Having obtained the abundance in the covered area, we can then scale-up to the study area:

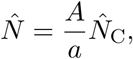

where A is the area of the study region to which to extrapolate the abundance estimate and *a* is the covered area. For line transects *a* = *2wL* (twice the truncation distance multiplied by the total length of transects surveyed, L) and for points *a* = *πw*^2^*T* (where *πw*^2^ is the area of a single surveyed circle and T is the sum of the number of visits to the sampled points).

We can use the Horvitz-Thompson-like estimator to calculate the “average” detectability for models which include covariates. We can consider what single detectability value would give the estimated 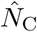 and therefore calculate:

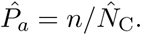

This can be a useful summary statistic, giving us an idea of how detectable our *n* observed animals would have to be to estimate the same 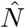 if there were no observed covariates. It can also be compared to similar estimates in mark-recapture studies. *P_a_* is included in the `summary` output and the table produced by `summarize_ds_models`.

*Stratification*

We may wish to calculate abundance estimates for some sub-regions of the study region, we call these areas *strata* and can be defined at the design stage or post hoc. Stratification can be used to increase the precision of estimates if we know *a priori* that density varies between different parts of the study area. For example, strata may be defined by habitat types which may be of interest for biological or management reasons. To calculate estimates for a given stratification each observation must occur in a stratum which must be labelled with a `Region.Label` and have a corresponding `Area`. Finally, we must also know the stratum in which each observation occurs.

As an example, the minke whale data contain two strata: `North` and `South` relating to strata further away from and nearer to the Antarctic ice edge, respectively.

### 5.2. Variance

We take an intuitive approach to uncertainty estimation, for a full derivation consult Marques and Buckland (2003). Uncertainty in 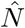 comes from two sources:

1. *Detection function*: Uncertainty in parameter (***θ***) estimation.
2. *Encounter rate*: Sampling variability arising from differences in the number of observations per transect.

We can see this by looking at the Horvitz-Thompson-like estimation in (1) and consider the terms which are random. These are: the detectability 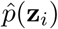 (and hence the parameters of the detection function it is derived from) and n, the number of observations.

Model parameter uncertainty can be addressed using standard maximum likelihood theory. We can invert the Hessian matrix of the likelihood to obtain a variance-covariance matrix. We can then pre- and post-multiply this by the derivatives of 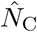 with respect to the parameters of the detection function

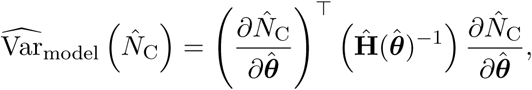

where the partial derivatives of 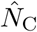 are evaluated at the MLE 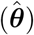 and H is the first partial Hessian (outer product of first derivatives of the log likelihood) for numerical stability (Buckland *et al*. 2001, p 62). Note that although we calculate uncertainty in 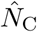, we can scale-up to variance of 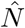 (by noting that 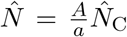 and therefore 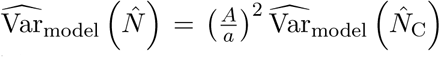, at least in the case where covariates are independent between strata (see e.g., Oedekoven, Buckland, Mackenzie, Evans, and Burger 2013).

*Encounter rate* is the number of objects per unit transect (rather than just n). When covariates are not included in the detection function we can define the encounter rate as *n/L* for line transects (where L is the total line length) or *n/T* for point transects (where T is the total number of visits summed over all points). When covariates are included in the detection function, it is recommended that we substitute the *n* in the encounter rate with the estimated abundance 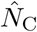 as this will take into account the effects of the covariates (Innes, Heide-Jørgensen, Laake, Laidre, Cleator, Richard, and Stewart 2002).

For line transects, by default, Distance uses a variation of the estimator “R2” from Fewster, Buckland, Burnham, Borchers, Jupp, Laake, and Thomas (2009) replacing number of observations per sample with the estimated abundance per sample, thus taking into account cluster size if this is recorded (Innes *et al*. 2002; Marques and Buckland 2003):

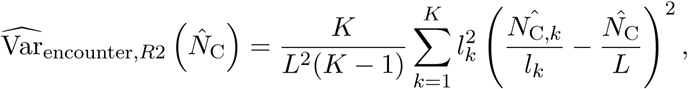

where *l_k_* are the lengths of the *K* transects (such that 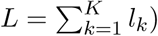 and 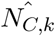 is the abundance in the covered area for transect *k*. For point transects we use the estimator “P3” from Fewster *et al*. (2009) but again replace n by 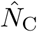 in the encounter rate definition, to obtain the following estimator:

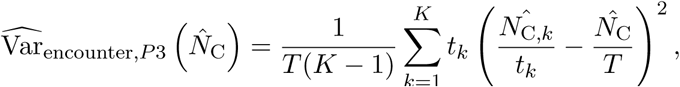

where *t_k_* is the number of visits to point *k* and 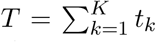 (the total number of visits to all points is the sum of the visits to each point). Again, it is straightforward to calculate the encounter rate variance for 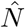 from the encounter rate variance for 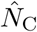.

Other formulations for the encounter rate variance are discussed in detail in Fewster *et al*. (2009). **Distance** implements all of the estimators of encounter rate variance given in that article. The `varn` manual page gives further advice and technical detail on encounter rate variance. For example for systematic survey designs, estimators S1, S2 and O1, O2 and O3 will typically provide smaller estimates of the variance.

We combine these two variances by noting that squared coefficients of variation (approximately) add (often referred to as “the delta method”; Seber 1982).

### 5.3. Estimating abundance and variance in R

Returning to the minke whale data, we have the necessary information to calculate *A* and *a* above, so we can estimate abundance and its variance. When we supply data to `ds` in the “flatfile” format given above, `ds` will automatically calculate abundance estimates based on the survey information in the data.

Having already fitted a model to the minke whale data, we can see the results of the abundance estimation by viewing the model summary:

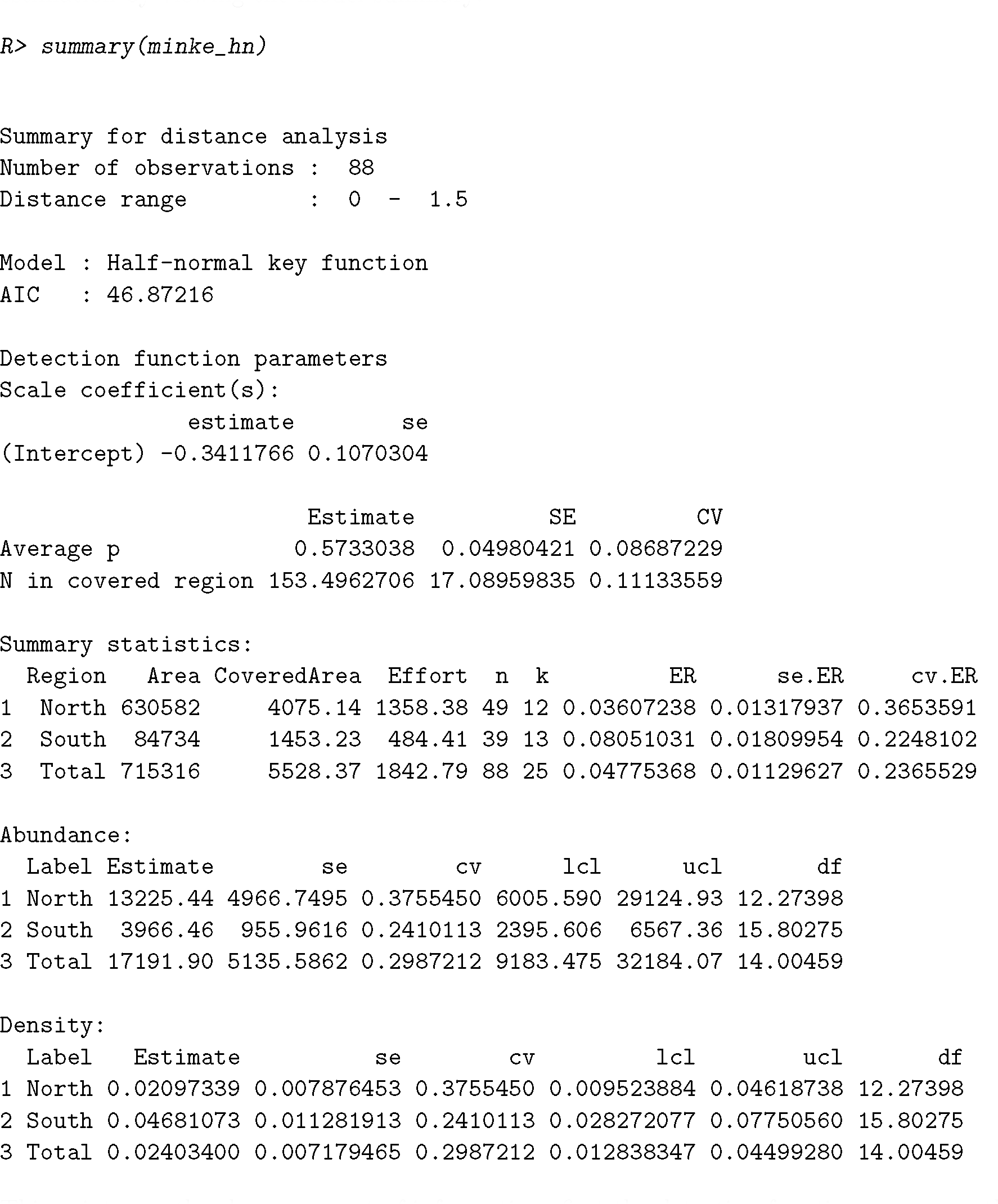

This prints a rather large amount of information: first the detection function summary, then three tables:

1. `Summary statistics`: giving the areas, covered areas, effort, number of observations, number of transects, encounter rate, its standard error and coefficient of variation for each stratum.
2. `Abundance`: giving estimates, standard errors, coefficients of variation, lower and upper confidence intervals and finally the degrees of freedom for each stratum’s abundance estimate.
3. `Density`: lists the same statistics as `Abundance` but for a density estimate.

In each table the bottom row gives a total for the whole study area.

The summary can be more concisely expressed by extracting information from the summary object. This object is a `list` of `data.frames`, so we can use the `kable` function from `knitr` to create summary tables of abundance estimates and measures of precision, such as Table 3. We prepare the `data.frame` as follows before using `kable`:

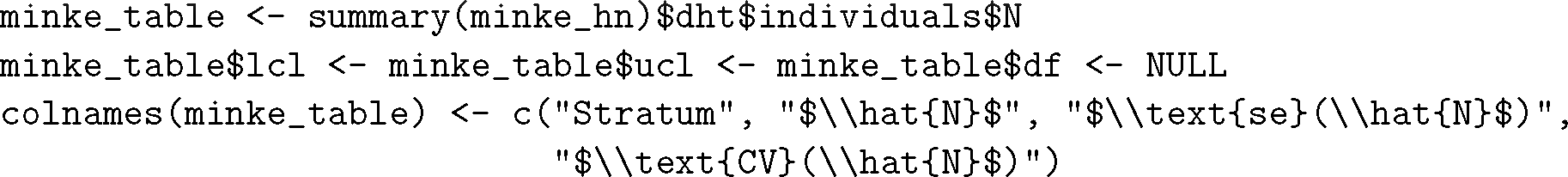

**Table 3.** Summary of abundance estimation for the half-normal model for the minke whale data.

| Stratum | $\hat{N}$ | $se(\hat{N})$ | $CV(\hat{N})$ |
| --- | --- | --- | --- |
| North | 13225.44 | 4966.750 | 0.376 |
| South | 3966.46 | 955.962 | 0.241 |
| Total | 17191.90 | 5135.586 | 0.299 |

## 6. Extensions

Distance sampling has been applied in a wide variety of situations. Objects being detected need not be animals; plants (Buckland, Borchers, Johnston, Henrys, and Marques 2007), beer cans (Otto and Pollock 1990), elephant dung (Nchanji and Plumptre 2001) and bricks on a lake bottom (Bergstedt and Anderson 1990) have been subjects of distance sampling investigations. The method of detection need not be visual sightings. Detections of objects can be through auditory as well as visual means (Marques, Thomas, Martin, Mellinger, Ward, Moretti, Harris, and Tyack 2013). Songs of birds (Buckland 2006) and whale vocalisations (Borchers, Pike, Gunnlaugsson, and Víkingsson 2009) are just two examples. Blows made by whales are examples of processes that can be modelled using distance sampling. Songs and blows are indirect sampling methods producing estimates of cue density. Cue densities can be converted to animal densities with additional data needed to estimate the rate at which cues are produced and the rate at which they disappear. These are but a few examples of the applications to which distance sampling has been applied (an incomplete list of references is given at http://distancesampling.org/dbib.html).

The features of **Distance** are deliberately limited to provide a simplified interface for users. For more complex analyses of distance sampling data, we provide additional packages for modelling in `R`.

We noted at the start of the article that **Distance** is a simple-to-use wrapper around the package **mrds**. Additional features available in **mrds** include models that relax the assumption that detection is certain at zero distance from the transect (by including data from additional observers). This is done using mark-recapture type methods which require additional survey methodology, known as double observer surveys or mark-recapture distance sampling (see Burt *et al*. 2014, for an introduction).

**Distance** can provide us with estimates of abundance or density for each strata as a whole but tells us nothing about the distribution of animals within strata. One option is to divide the study area into smaller and smaller strata to try to detect patterns in spatial distribution, however, a more rigorous approach is to build a spatial model. Such models incorporate spatially-referenced environmental data (for example derived from GIS products). **Distance** interfaces with one such package used to perform this type of analysis: **dsm** (Miller, Rexstad, Burt, Bravington, and Hedley 2015). So-called “density surface modelling” uses the generalized additive model framework (e.g. Wood 2006) to build models of abundance (adjusting counts for imperfect detectability) as a function of environmental covariates, as part of a two stage model (Hedley and Buckland 2004; Miller, Burt, Rexstad, and Thomas 2013).

Uncertainty in measured covariates (e.g. cluster size) and model uncertainty (when two models have similar fit but substantially different estimates) can be incorporated using the multi-analysis distance sampling package **mads** (Marshall 2015b). In addition, **mads** can also incorporate sightings with unknown species identification. This is done by estimating the abundance of these unidentified sightings and pro-rating them to the known species (Gerrodette and Forcada 2005).

As mentioned above, survey design is critical to ensuring that resulting distance sampling data can be reliably analysed. **DSsim** allows users to test out different designs in their study region and tailor population attributes to reflect the species they are working with. **DSsim** (Marshall 2015a) allows users to more easily identify challenges unique to their study and select a survey design which is more likely to yield accurate and precise estimates.

Distance for Windows has many users (over 50,000 downloads since 2002) and they may be overwhelmed by the prospect of switching existing analyses to `R`. For that reason we have created the **readdst** (Miller 2015) package to interface with projects created by Distance for Windows. The package can take analyses created using the CDS, MCDS and MRDS engines in Distance for Windows, extract the data and create equivalent models in `R`. **readdst** can also run these analyses and test the resulting statistics (for example, 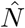 or 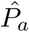) calculated in `R` against those calculated by Distance for Windows. We hope that **readdst** will provide a useful transition to `R` for interested users. **readdst** is currently available on GitHub at https://github.com/distancedevelopment/readdst.

## 7. Conclusion

We have given an introduction as to how to perform a distance sampling analysis in `R`. We have covered the possible models for detectability, model checking and selection and finally abundance and variance estimation.

In combination with tools such as **knitr** and **rmarkdown** (Allaire, Cheng, Xie, McPherson, Chang, Allen, Wickham, Atkins, and Hyndman 2015), the helper functions in **Distance** provide a useful set of tools to perform reproducible analyses of wildlife abundance for both managers and ecologists. We also direct readers’ attention to **DsShiny** (Laake 2014), a package that builds on **Shiny** (Chang, Cheng, Allaire, Xie, and McPherson 2016) and **mrds** to allow users to build and fit models in a graphical interface, with output to RMarkdown.

**R** and its extension packages provide many tools exploratory data analysis that can be useful for a distance sampling analysis. We hope that this paper provides useful examples for those wishing to pursue distance sampling in `R`. More information on distance sampling can be found at http://distancesampling.org and a mailing list is maintained at https://groups.google.com/forum/#!forum/distance-sampling.

We note that there are other packages available for performing distance sampling analyses in `R` but believe that **Distance** is the most flexible and feature-complete, and provides pathways to a range of more complicated analyses. Appendix A gives a feature comparison between **Distance** and other `R` packages for analysis of distance sampling data.

## 8. Acknowledgements

The authors thank the two anonymous reviewers for their constructive comments, which have greatly improved the paper and package. The authors would like to thank the many users of **Distance, mrds** and Distance for Windows who have contributed bug reports and suggestions for improvements over the years. We would particularly like to thank Steve Buckland, David Borchers, Tiago Marques, Jon Bishop and Lorenzo Milazzo for their contributions. We thank Tiago Marques a second time for helpful comments on an early version of the paper and Colin Beale who suggested the “flatfile” data format. We also thank Ken Burnham and David Anderson for fundamental contributions to the early development of these methods.

## Appendix A: Feature comparison

There are four packages available for analysis of distance sampling data in `R` that we are aware of. All are available on CRAN. On top of **Distance** and **mrds** (Laake *et al*. 2015) described above, they are **Rdistance** (McDonald, Nielson, and Carlisle 2015) and **unmarked** (Fiske and Chandler 2011). Table 4 provides a feature comparison of these packages.

### Abundance estimation

**Distance** and **mrds** models can be used as part of a density surface model using **dsm**, which allows abundance to be modelled as a function of spatially varying covariates (such as location, sea depth, altitude etc). See Miller *et al*. (2013) for more information.

**unmarked** also allows abundance to vary according to covariates, via the abundance part of the likelihood. See Fiske and Chandler (2011) for more information on the package and Royle, Dawson, and Bates (2004) for more information on methodology.

Resulting **Rdistance** models can be use in combination with `R` modelling functions such as `lm, glm` etc to build abundance estimates which vary according to covariates. More information is available on the project’s wiki.

**Table 4.** Feature comparison of the available packages to perform distance sampling analyses.

| Feature | Distance | Rdistance | unmarked | mrds |
| --- | --- | --- | --- | --- |
| Line transects | x | x | x | x |
| Point transects | x |  | x | x |
| Interval (binned) distances | x |  | x | x |
| Exact distances | x | x |  | x |
| Continuous individual level covariates | x |  |  | x |
| Factor individual level covariates | x | x |  | x |
| Transect level covariates | x |  | x | x |
| Objects in clusters | x | x | x | x |
| Left truncation | x | x |  | x |
| Half-normal key | x | x | x | x |
| Hazard-rate key | x | x | x | x |
| Uniform key | x | x | x | x |
| Gamma key |  | x |  | x |
| Negative exponential key |  |  | x |  |
| Adjustment terms | x | x |  | x |
| AIC adjustment selection | x | x |  |  |
| Monotonicity constraints | x |  |  | x |
| Availability bias model |  |  | x |  |
| Perception bias model |  |  |  | x |
| Abundance estimation | x | x | x | x |
| Density estimation | x | x | x | x |
| User-defined likelihood functions |  | x |  |  |

